# ExtendAlign: the post-analysis tool to correct and improve the alignment of dissimilar short sequences

**DOI:** 10.1101/475707

**Authors:** Mariana Flores-Torres, Laura Gómez-Romero, Joshua I. Haase-Hernández, Israel Aguilar-Ordóñez, Hugo Tovar, S. Eréndira Avendaño-Vázquez, C. Fabián Flores-Jasso

## Abstract

In this work, we evaluated several tools used for the alignment of short sequences and found that most aligners execute reasonably well for identical sequences, whereas a variety of alignment errors emerge for dissimilar ones. Since alignments are essential in computational biology, we developed ExtendAlign, a post-analysis tool that corrects these errors and improves the alignment of dissimilar short sequences. We used simulated and biological data to show that ExtendAlign outperforms the other aligners in most metrics tested. ExtendAlign is useful for pinpointing the identity percentage for alignments of short sequences in the range of ∼35–50% similarity.

## BACKGROUND

Since the emergence of high-throughput sequencing technologies, the development of computational tools that aid in the assembling, comparison, and analysis of multiple genomic sequences has flourished tremendously [1,2]. The sequencing of genomes from novel organisms has led to important discoveries that have shaped our understanding of what makes species alike, and also unique, at the genomic level [2–6]. The analysis of gene expression by high-throughput sequencing has become the workhorse for many laboratories seeking to understand how cells respond to biochemical and metabolic changes or external stimuli [3,4,7,8]. For all those studies for which there exist a corresponding reference genome, one of the first steps of the computational analysis is the alignment of sequencing reads to its reference [3,9]; for those for which there is no corresponding reference yet, it is common to employ a related genome for the alignment of reads [9–11]. Although this strategy has led to important discoveries over the last decade about the expression profile, mutation landscape, or sample diversity, for species whose genomes are still unknown, the computational analysis has been constrained primarily to those sequences for which there is a high similarity level compared to their references [10–14]. For dissimilar sequences, however, it is more challenging to assign or infer the biological context for which they function [11,12], and therefore it is also common to set those apart until there is an appropriate cognate reference.

Although sequencing technologies have evolved to yield longer sequencing reads compared to the early beginnings of the genomic era, the vast majority of studies rely on the use of sequencing platforms whose reads range ∼50–150 nt in length [1,3,4]. There are a number of computational tools based primarily on two main types of pairwise algorithms which constitute the current and most popular methods for the alignment of sequences within this range: local and global algorithms [15–17]. Local alignments identify similar regions in sequences by determining homology in the presence of rearrangements [18–21]. Global alignments find similarity by transforming the aligned sequences into one another by a combination of simple edits [5,6,22–24]. While most aligners based on these two approaches execute reasonably well for alignments of identical, or nearly identical sequences, their accuracy and robustness decrease considerably as the similarity between sequences declines, particularly for sequences shorter than 30 nt.

In this work, we explored how the dissimilarity affects the alignment of short sequences by comparing the results of several computational tools widely used for the alignment of short sequences. We find that, while most aligners perform reasonably well with identical or nearly identical short sequences, they execute poorly in the alignment of dissimilar ones. Depending on the underlying algorithm, the aligners compared retrieve alignments that differ in the type of errors produced: local aligners tend to clip nucleotides flanking the seed substring, which results in alignment reports that miss matches or mismatches; global aligners do not fail in reporting matches, but introduce gaps to achieve end-to-end alignments, which results primarily in a low precision score.

The accumulation of these errors sets a challenge for the computational analysis of short sequences and has an impact on all those alignments for which there is no corresponding reference available [10,12]. For example, studies aimed at discovering small RNAs from novel and distant species are inevitably forced to employ genome references other than their own — if the dissimilarity between sequences is extensive, the alignment results may bias the interpretation.

To address this issue, we developed ExtendAlign, a post-analysis tool that provides a significant improvement for the alignment of dissimilar short sequences. ExtendAlign quantifies the identity percentage in the alignment based on the accurate number of matches and mismatches that may initially be missed by a local alignment. Since it incorporates the output of a local algorithm and provides an end-to-end alignment report, ExtendAlign combines the strength of a multi-hit local alignment, with the refinement provided by a query-based global algorithm without applying a clipping strategy. Therefore, its reports position at the intersection between local and global alignments for short sequences. We evaluated the performance of ExtendAlign with simulated and biological data and show that it outperforms other computational tools commonly employed to align short sequences in most of the metrics tested.

By executing multiple alignments with short sequences against distant genomes as references, we show that ExtendAlign is particularly useful to recalculate and pinpoint the identity percentage of alignments that span across the “twilight zone” —the similarity that ranges ∼35–50% [25,26]. Finally, we provide one practical example of the utility of ExtendAlign by revisiting the alignment of RNA sequences considered to have a bovine-specific origin contained within library datasets obtained from human samples of published literature —a recent biological controversy raised by the report of contaminating RNAs from cow into cell lines due to their culture with fetal bovine serum [27,28]. We found that more than 35% of all small RNA sequences considered bovine-specific present in human cell lines are at least 80% identical to humans. This indicates that — because of the aligner and specific parameters employed— there was a high false-positive discovery rate of bovine-specific small RNAs that contributed to this controversy.

ExtendAlign is recommended for short sequence alignments that require the highest accuracy; or for studies that require a quantitative measure of the dissimilarity level; or for studies where precision in the identity percent cut-off is critical for determining homology between phylogenetically distant short sequences. ExtendAlign was developed as a Nextflow pipeline to guarantee reproducibility and scalability [29], and it is available for download at https://github.com/Flores-JassoLab/ExtendAlign.

## RESULTS

### Sequence dissimilarity impacts the alignment of short sequences

We first examined the results provided by tools commonly employed for the alignment of short sequences to get an insight into their alignment capabilities. Two identical sequences were aligned with Bowtie, Bowtie2, BWA, BWA-MEM, BLASTn, BLASTn-short, and Needle (Additonal file 1: Figure S1A, left) [18,22,30–33]. Except for BWA-MEM, all aligners retrieved a full-length alignment hit under default parameters. This simple comparison suggests that most of these aligners can handle short sequences if the purpose is the identification of perfect or near-perfect alignments; which is often the case for reads from high-throughput sequencing samples of small RNAs to a related reference genome. However, when two dissimilar sequences were aligned, only Needle and BLASTn-short retrieved alignment hits under default parameters (Suppl. Figure S1A, right). Needle is an aligner based on the Needleman-Wunch algorithm and performs end-to-end alignments [22]. The Bowtie, BWA and BLASTn versions tested are based on local algorithms, and therefore require a seed substring of a minimum fixed size to initiate alignments. The algorithmic approaches employed by Needle and BLASTn-short retrieve different alignments for the same two sequences under default parameters (Suppl. Figure S1B). For example, Needle managed to find the positions matched correctly, but reported overhanging positions as gaps.

The fact that only BLASTn-short retrieved alignment results compared to the other local algorithms motivated us to examine in more detail whether this absence of hits was due to the general intrinsic capabilities of local alignments. Hence, we adjusted the parameters of every tool tested to make all seed sizes uniform, and also to maximize their alignment ability —which is concomitantly associated with an increase in the alignment hits (see Suppl. Table S1). Under these permissive parameters, BWA-MEM retrieved an alignment hit (Suppl. Figure S1A, right), albeit different to those of BLASTn and BLASTn-short (Suppl. Figure S1C); Bowtie, Bowtie2, and BWA did not retrieve alignments. Interestingly, regardless of the parameters employed, there were positions with identical nucleotides not reported primarily outside the seed substring and near the 5’ or 3’ ends in the query (Suppl. Figure S1B and S1C, bold red letters). We rule out that the permissive parameters chosen prevented the local aligners from performing efficiently since they all found a full-length alignment for the identical sequences (Suppl. Figure S1A, left); arguing in favor of their popularity for aligning sequences this short, despite the recommended use by their developers (Suppl. Table S1) [18,22,30–33].

To investigate further how sequence dissimilarity impacts the alignment of short sequences, we simulated query and subject databases and aligned them with all the aligners mentioned above. Inaccuracies in the alignment of short sequences might affect the study of several classes of RNAs, for example, small RNAs [34]. An abundant class of small RNAs are microRNAs, and since their mature sequence sizes peak at ∼22 nt in plants and animals [35–38], the simulated query database consisted of 8,500 sequences randomly generated of 22 nt long. The subject database consisted of sequence sizes ranging from 50–170 nucleotides with fixed 7mer seed, each with a randomly chosen position 1, and one or up to fifteen mismatches, insertions or deletions (see Methods for details).

Under default parameters, all aligners showed a balanced performance among the metrics tested (except for BWA-MEM, which did not retrieve alignment hits) (Table 1). For example, BLASTn-short retrieved the largest number of hits, at the cost of Sensitivity and Specificity; Needle retrieved the highest number of True Positive Hits and the highest Recovery Rate (7,038 and 0.83, respectively), but was also the least precise of all tools tested; the Bowtie versions tested and BWA retrieved a small number of alignment hits, and their Recovery Rate was also low but showed high Sensitivity compared to the other aligners. Importantly, Needle retrieved an equal number of Hits to queries, while the BLAST versions exceeded this number, reflecting that there was more than one alignment hit per each query. Under permissive parameters, in contrast, all local aligners increased the number of Hits, being Bowtie2 the highest. Both BLASTn and BLASTn-short retrieved the highest number of True Positives, as well as Recovery Rate and Precision (7,202, 0.85, and 0.98, respectively); albeit also presented the least Sensitivity and Specificity (0.26 and 0.001, respectively). BWA-MEM showed a drastic improvement in several metrics, indicating that a parameter adjustment might have a severe impact on sequence alignments.

Overall, the results of this simulation imply that there are fundamental limitations for the alignment of dissimilar short sequences with current methods —while some metrics performance improved under specific parameters, others decrease; regardless of the parameters, no aligner excelled in all the metrics tested. Primarily, the local approaches miss identical nucleotides near the 5’ or 3’ ends on the alignment, or provide erroneous reports on the number of matches, particularly for dissimilar sequences with several gaps or mismatches flanking the seed substring (Suppl. Figure S1B and S1C, and data not shown). Needle, on the contrary, showed the least precision of all aligners because its alignment report is based on the farthest 5’ and 3’ ends within each alignment (Suppl. Figure S2A and S2B, and data not shown). The cumulative errors caused by either algorithm might give rise to alignment biases, particularly while aligning thousands of dissimilar sequences to establish similarity between short sequences obtained from high-throughput sequencing data. Compromising the alignment accuracy of short sequences may have an impact on the understanding of small RNAs; for example, by restraining the identification of evolutionary relationships that might exist among microRNAs of distant species [12,38].

To overcome this problem, we developed ExtendAlign, a post-analysis tool that improves the alignment results of dissimilar short sequences by correcting the errors mentioned above (Figure 1A). ExtendAlign identifies the number and identity of all nucleotides flanking a seed substring in a query that might have gone unaligned. After finding unreported nucleotides, ExtendAlign recalculates and reports the total number of matches and mismatches (m/mm) in an end-to-end manner for each query. Therefore, alignment biases are diminished by: i) accounting for all undetected m/mm, ii) considering overhanging nucleotides in the query as mismatches, and iii) extending the alignment to the 5’ and 3’ ends in the query (not the subject). Common examples of alignment errors and their corrections are shown in Figure 1B. Because of its low sensitivity and specificity, but also because of its high recovery rate and precision under default and permissive parameters, ExtendAlign was developed to function after priming the alignments with the BLASTn versions tested in this work. Importantly, BLASTn-short is a version of BLASTn aimed to align short sequences [33,39]; however, the permissive parameters of BLASTn-short and BLASTn of this work are equivalent, and hereinafter are referred to as *high sensitivity*-BLASTn (HSe-BLASTn).

**Figure 1.**
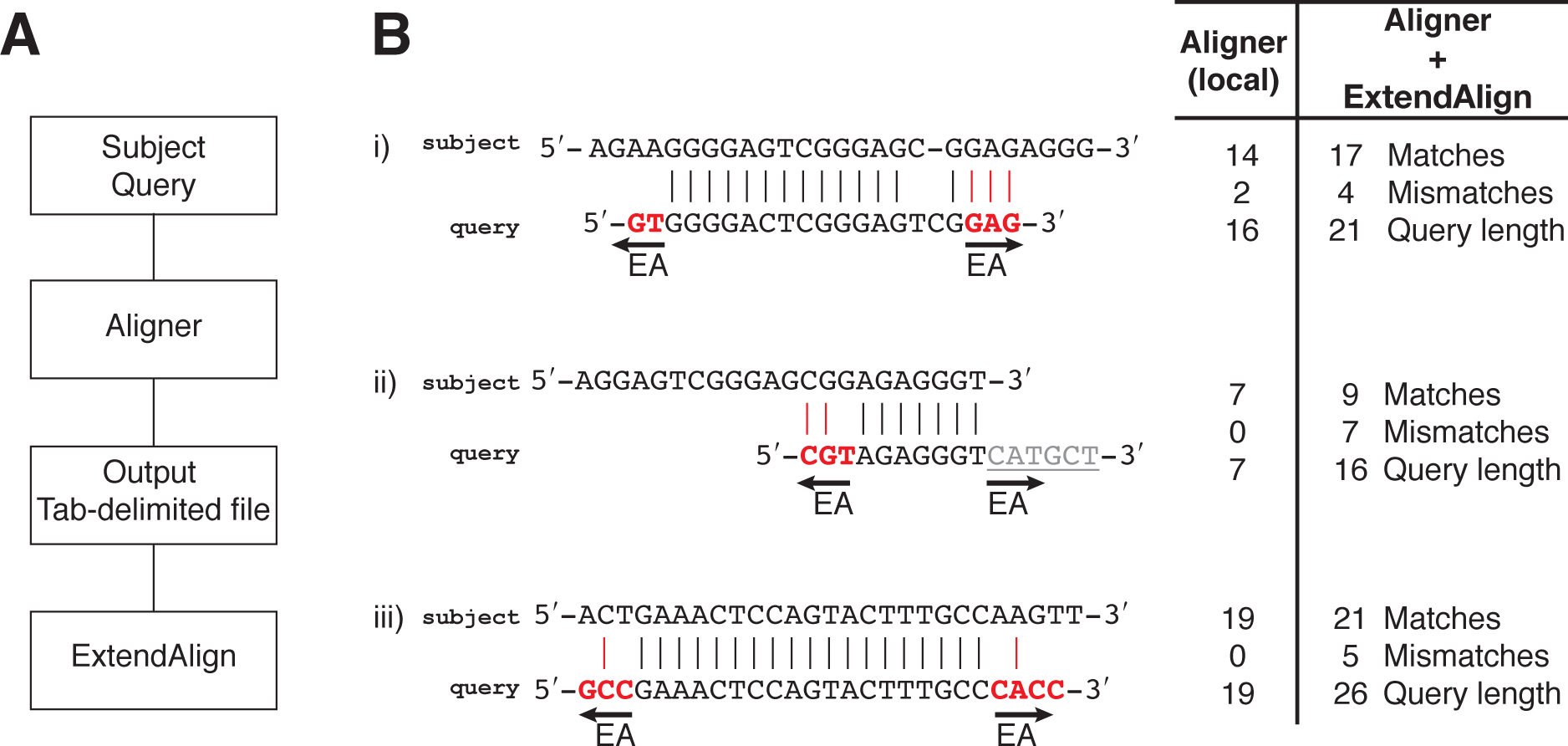
ExtendAlign is a post-analysis tool to correct for errors created during the alignment of dissimilar short sequences. A, ExtendAlign (EA) is designed to incorporate the output of a local pairwise aligner as tab-delimited file and improve the alignment results. B, Examples of unreported nucleotides by pairwise local aligners comprising matches or mismatches that originate alignment biases and the correction that takes place after EA. Unreported nucleotides (bold red letters) are identified based on the query length and extended to identify possible m/mm (i–iii). Overhanging positions are reported as mismatches only for the query (ii, underlined grey letters); alignments are based solely on the query length.

### ExtendAlign increases the sensitivity and specificity of local alignments

We examined the performance of ExtendAlign using the previous simulated databases and measured all the metrics tested (Table 2). In general, ExtendAlign showed an improved or similar performance in most metrics compared to the other aligners tested under default and permissive parameters. For example, Sensitivity and Specificity increased more than 3-fold and 30-fold, respectively, compared to HSe-BLASTn; which is comparable with the performance score of other aligners. Despite being the highest-ranked compared to other aligners, Precision was the only metric for which ExtendAlign did not improve in comparison to its priming base. We attribute this result to how the alignment correction takes place: ExtendAlign identifies nucleotides flanking the seed substring that might have gone unnoticed by the local alignments (Figure 1B); thus, the apparent lower Precision score might be a consequence of increasing the alignment length and also decreasing the number of False Negatives.

Next, we analyzed how the alignment correction by ExtendAlign compares to local and global approaches. For this, we chose HSe-BLASTn and Needle as they provide the highest Number of True Positive Hits in our simulation analysis. Since Needle alignments are paired-wised, only 8,500 hits were retrieved, whereas HSe-BLASTn retrieved 4,715,796 hits due to its multiple-hit capabilities (Table 1 and Suppl. Figure S3A). Hence, to compare the results for each alignment approach, all the alignment pairs located by HSe-BLASTn with one-hit only were presented to Needle (4,561,877 pairs) (Suppl. Figure S3B, and Suppl. Info). Finally, the total number of hits for each aligner was plotted as a function of sequence coverage and total alignment size, gaps, or mismatches (Figure 2). HSe-BLASTn retrieved a variety of query sequence coverage that ranged from ∼30–100%, and whose total alignment length ranged from 7–22 nt, with a maximum of four gaps, and up to six mismatches per query (Figure 2A–C). Needle, on the contrary, retrieved 100% coverage for every alignment pair, but the alignment size and the number of gaps were raised to 150 nt (Figure 2A and 2B). This implies that Needle: i) considers overhanging nucleotides in the query or subject as part of the alignment, and ii) introduces gaps interspersed along some sequences to achieve alignments (Suppl. Figure S2A and S2B). This result makes Needle impractical for aligning sequences that vary in size, for instance, finding multiple loci for short sequences against whole chromosomes or genomes, which would result in a different type of bias in comparison with local algorithms (Suppl. Figure S3A). Conversely, ExtendAlign does not increase the total alignment size, nor introduces gaps, and most importantly, since it always achieves 100% query coverage, does not miss mismatches flanking the seed substring. Due to these results, we conclude that ExtendAlign is a robust post-analysis tool that refines the alignment of dissimilar short sequences and provides exceptional accuracy because of its end-to-end correction capability. The features and recommended use of ExtendAlign are listed in the Supplementary Table S1.

**Figure 2.**
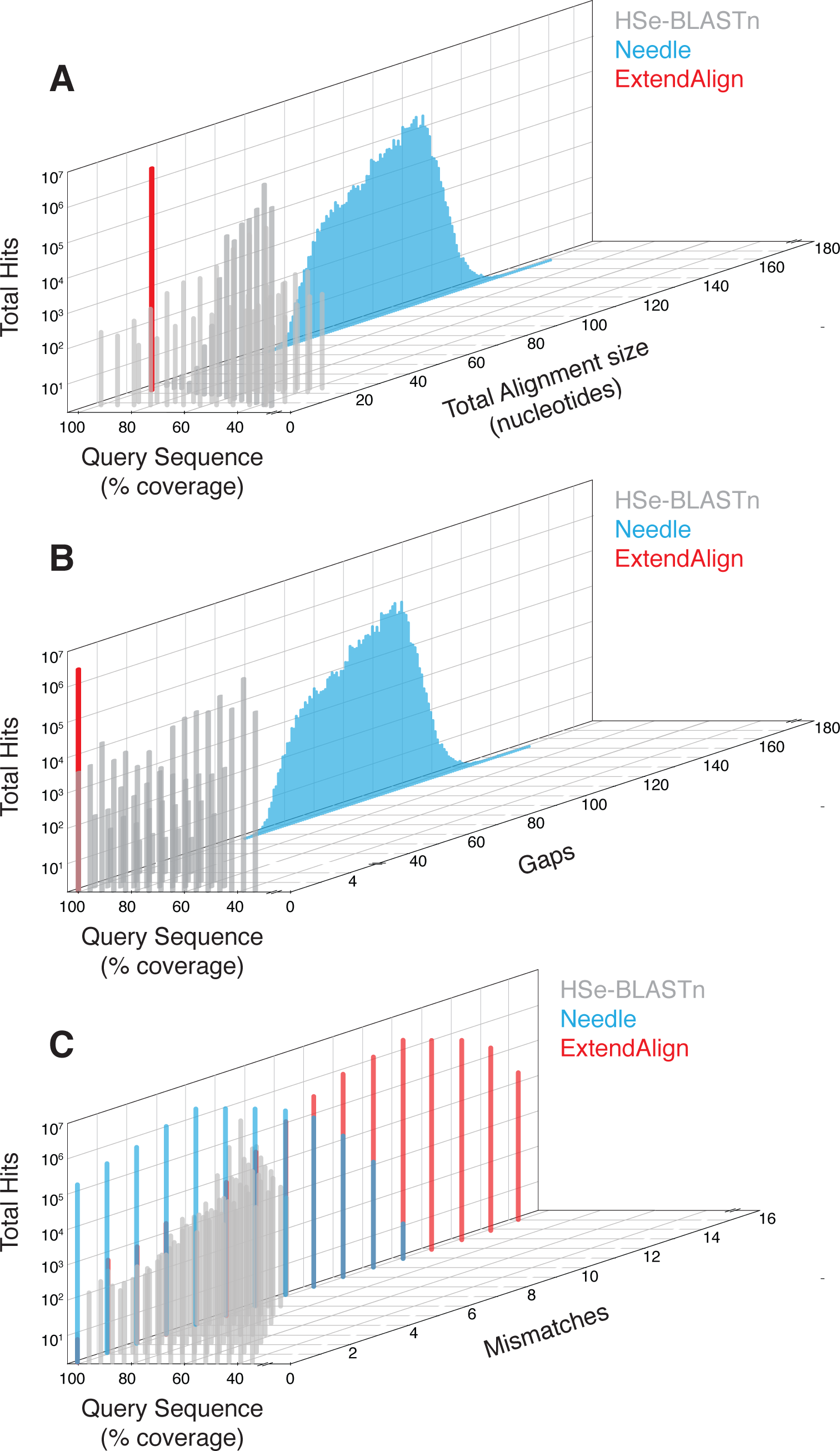
ExtendAlign correction positions at the intersection between local and global approaches. A–C, The one-hit per alignment sequence pairs obtained with HSe-BLASTn (grey) from the simulated databases analysis were presented to Needle (cyan), and compared to ExtendAlign (red). The coverage percent of query aligned was plotted as a function of the total number of hits for the total alignment size (A), the number of gaps (B), and the number of mismatches per alignment (C).

### ExtendAlign increases the number of total matches and mismatches for dissimilar alignments

We further evaluated ExtendAlign to get a better understanding of its precision. If increasing the alignment coverage impacts the Precision score, it should be reflected directly in a concomitant increase in the number of m/mm identified, presumably at positions located near the 5’ or 3’ ends. In a biological context, it is expected to find multiple examples of dissimilar alignments if different short sequences are aligned to a large reference, for example, a whole genome. Hence, we aligned all human microRNAs against the mouse genome and measured directly the number of m/mm for all alignments. Being conserved among mammals, the mouse genome contains several identical loci to most human microRNAs [40], but also a plethora of dissimilar loci to look for m/mm. After the alignments of microRNAs against a related genome reference, however, we observed only a marginal increase in the total number of m/mm after the correction by ExtendAlign compared to those of HSe-BLASTn (Figures 3A and 3B, main graphs, *p* < 0.0001; Kolmogorov-Smirnov *D* = 0.005, matches; and *D* = 0.016, mismatches).

**Figure 3.**
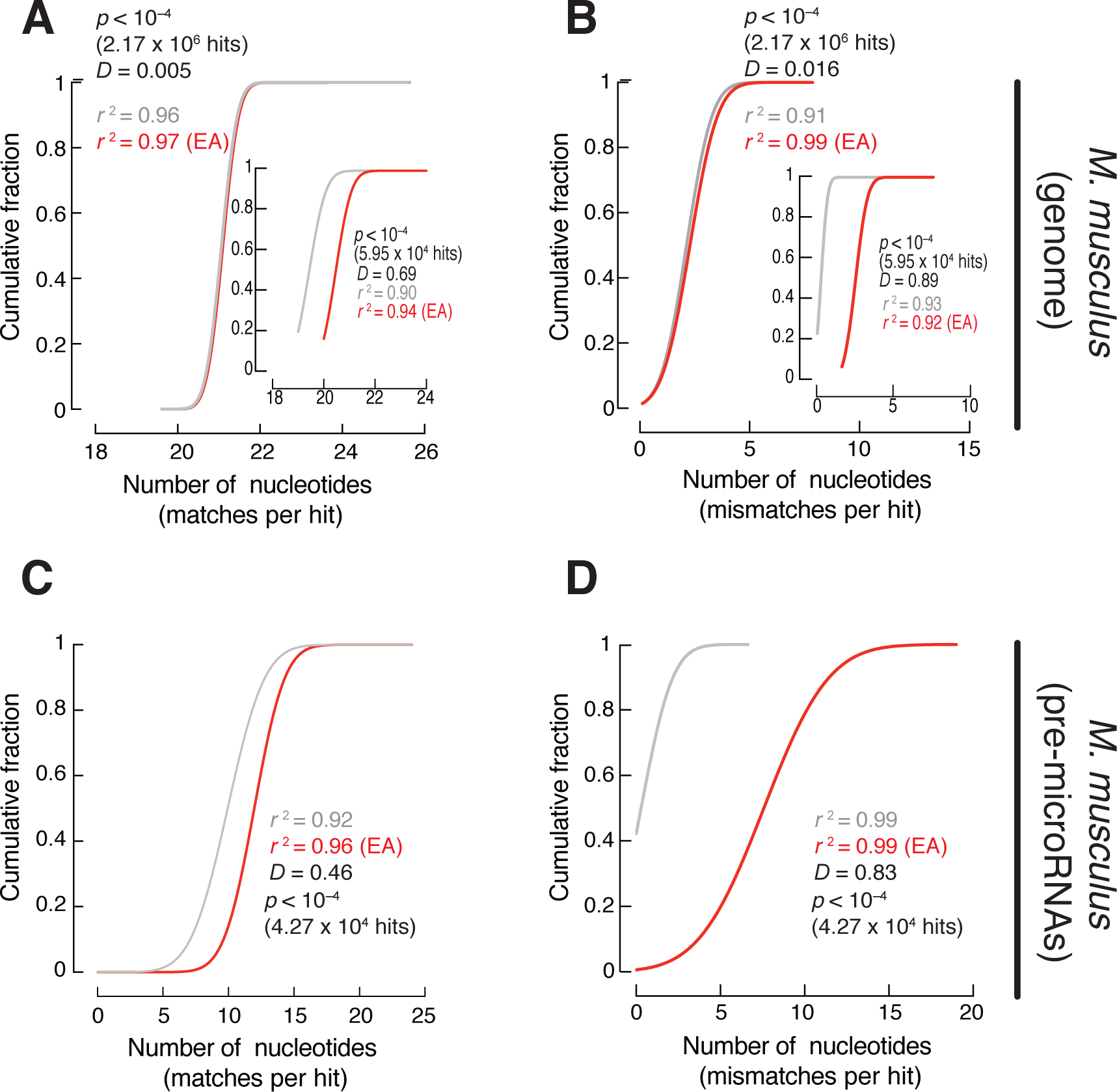
ExtendAlign improves alignments by increasing significantly the number of matches and mismatches. The total number of matches (A and C), and mismatches per hit (B and D), were plotted as a cumulative fraction for the alignments of human mature microRNAs against the mouse genome (A and B), or against pre-microRNAs (C and D). Insets in A and B correspond to the number of m/mm from alignments whose perfect-matched hits were removed. Grey, HSe-BLASTn; red, ExtendAlign. Significance values were calculated using the Kolmogorov-Smirnov test (K-S) for unpaired nonparametric data; *D* = K-S distance. The non-linear fit curve by least squares as used as measure of goodness of fit.

One explanation for this result is that the amount of perfect, or near-perfect loci spotted by HSe-BLASTn alone, vastly outnumbers the net correction by ExtendAlign (Suppl. Table S2). This is further supported by the results obtained when mature microRNAs were aligned only against the mouse precursor microRNAs (Figure 3C and 3D, Kolmogorov-Smirnov *D* = 0.46, matches; and *D* = 0.83, mismatches; *p* < 0.0001). Precursor microRNAs (pre-microRNAs) size range is ∼60–75 nt [41,42]. Consequently, an eight-fold increase in the total number of mismatches imply that their alignment to mature sequences largely constrain the chance of finding perfect match hits compared to an entire genome that could mask the correction with ExtendAlign (Suppl. Table S2, mismatches). Therefore, since perfect-match hits are not candidates for correction, we removed them from the m/mm counts. Accordingly, the correction showed a markedly less marginal difference compared to HSe-BLASTn, for both matches and mismatches (Figures 3A and 3B, inset graphs; *p* < 0.0001).

Regardless of the subject reference or the exclusion of perfect-matched hits, we noticed a more drastic difference between the total number of matches *vs*. mismatches in all cases —*e.g*., microRNAs against genome: *D* = 0.005 *vs. D* = 0.016, and *D* = 0.69 *vs. D* = 0.89; microRNAs against pre-microRNAs: *D* = 0.46 *vs. D* = 0.83. This strongly suggested that the correction by ExtendAlign is more effective if more divergent sequences are aligned because mismatches are a direct measurement of similarity [9,43]. For this reason, we tested whether the alignment of short sequences against less conserved genomes showed similar behavior. For this, we used as examples the genome of *Dasypus novemcinctus* (armadillo) because it is at the border of taxonomic classification, and due to the large content of dissimilar sequences in its genes it is difficult to assign the correct taxa [44]; and also the *Ornithorhynchus anatinus* (platypus) genome because it diverged from the mammalian lineage 160 million years ago, and thus possesses a unique blend of morphological and genomic features of mammals, reptiles, birds and fish [45]. The alignment of human pre-microRNAs revealed a significant increase in the total number of m/mm for both genomes after the ExtendAlign correction (Figure 4A–D). Importantly, in contrast to the alignment results observed for mouse genome (Figure 3A and 3B), there is a noticeable difference in the total m/mm compared to HSe-BLASTn in all cases without the need of removing perfect-matched sequences from the alignment, further supporting the utility of ExtendAlign to align dissimilar short sequences. The difference observed is also more drastic in matches compared to mismatches (armadillo: *D* = 0.43 *vs. D* = 0.91; platypus: *D* = 0.12 *vs. D* = 0.36). Since Precision is calculated based on the number of mismatches (true and false positives), their increase has a direct effect on this metric. Taking together all these results, we conclude that the cumulative bias observed in the alignment of dissimilar short sequences mainly results from mismatched nucleotides not covered during the local alignment; identifying such missing positions reflects therefore an apparent decrease in the precision score.

**Figure 4.**
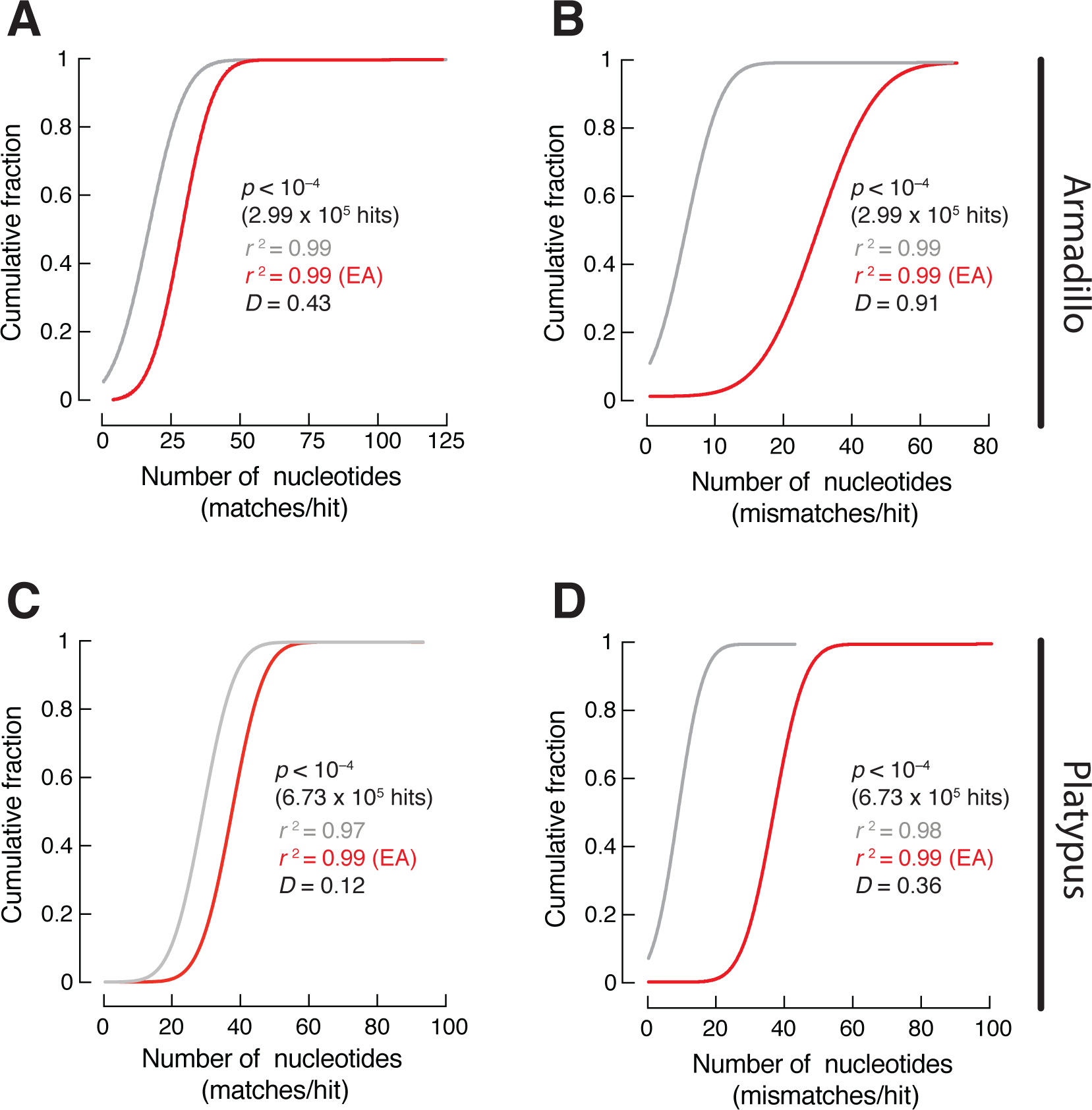
Dissimilar alignments are suitable targets for correction. The total number of matches (A and C), and mismatches per hit (B and D), were plotted as a cumulative fraction for the alignments of human pre-microRNAs against the armadillo genome (A and B), or against the platypus genome (C and D). Grey, HSe-BLASTn; red, ExtendAlign. Significance values, K-S distance, and goodness of fit, were measured as in Figure 3.

### Pinpointing the similarity percentage of short sequences with ExtendAlign

Next, we evaluated the utility of ExtendAlign to classify short sequences according to their similarity percent against a reference. The lack of similarity between distant short sequences has an impact on predicting secondary structures in nucleic acids [25,26]. As the similarity between sequences decreases and approaches the so-called “twilight zone” —the similarity that ranges from ∼35–50% identity, it is generally assumed that secondary structure predictions might not be reliable because alignments at this range tend to obscure the covariance signal [25,26,46]. From the alignments of pre-microRNAs *vs*. the armadillo and platypus genomes, we classified all alignment hits according to their identity percentage to each reference. Fifty percent of all hits achieved by HSe-BLASTn to the armadillo and platypus genomes span across this range, while 25% of hits reached ∼35% or less similarity (Figure 5A and 5B). After the ExtendAlign correction, the median identity to each reference increased from 43% (armadillo) and 41% (platypus), to more than 50% each (*p* < 0.0001). The increase above the twilight zone of half the hits indicates an improvement in the alignment of dissimilar short sequences.

**Figure 5.**
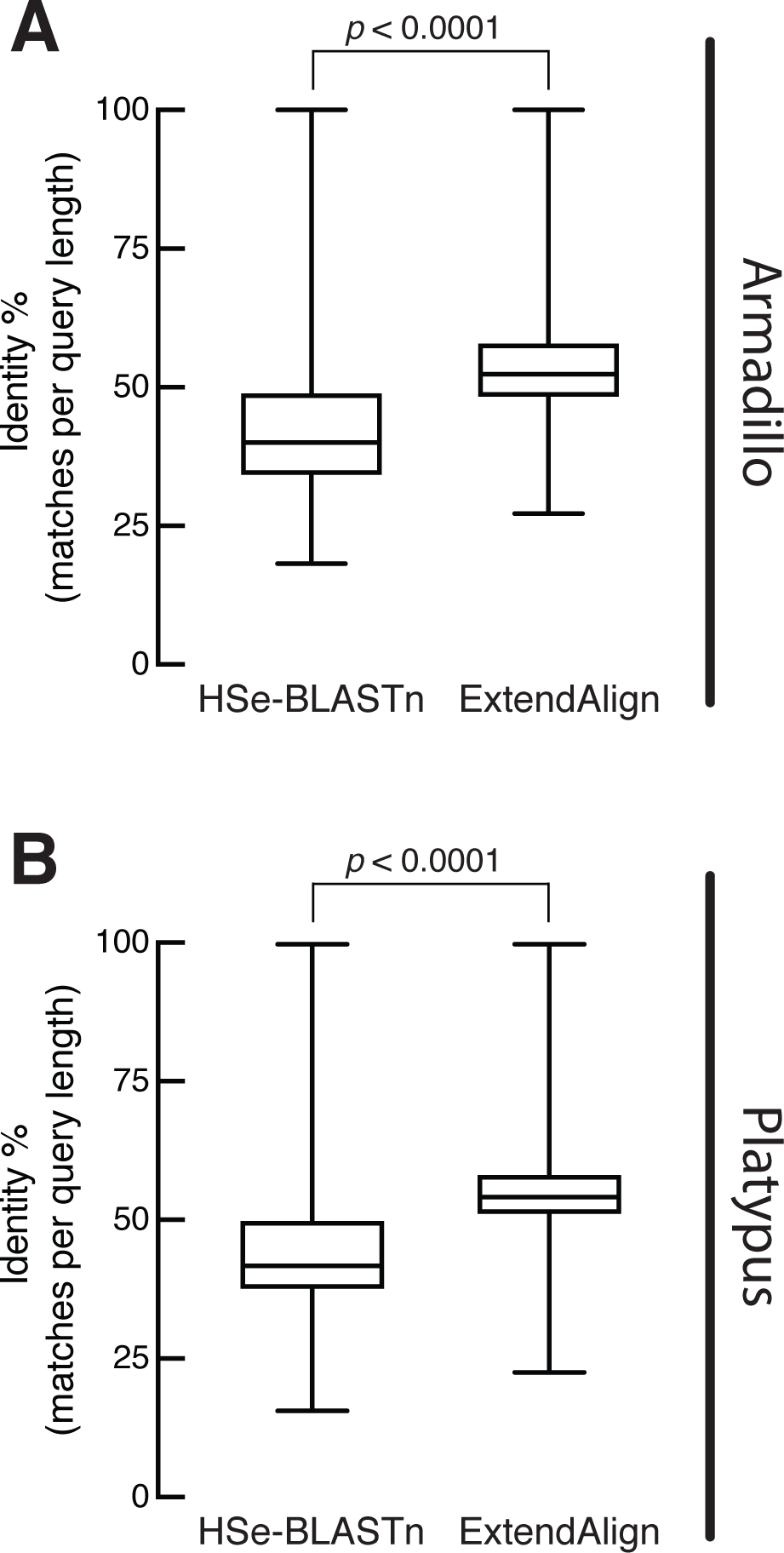
ExtendAlign is useful to correct alignments that span across the twilight zone. The identity percentage of all human pre-microRNAs was measured as matches per query length and compared for HSe-BASTn and ExtendAlign. A, armadillo genome; B, platypus genome. Box plots are min to max values shown by quartiles; significance was obtained by using the Wilcoxon test for paired, non-parametric data.

Finally, we used ExtendAlign to revisit the alignments performed in a recent work that reported that small RNA contained in fetal bovine serum (FBS) might transfer into cell cultures [27]. The extent at which this observation might affect the entire literature about microRNAs function is unknown, and still a matter of debate [28]. In their work, Wei and colleagues performed a computational analysis that encompassed the search for “bovine-specific” RNA sequences contained within public sequence read archives (SRAs) from human samples. By using Bowtie2, the authors considered all the sequences that aligned to the cow genome, but did not to the human genome, as bovine-specific (Figure 6A). Since the analysis was made using SRAs aimed to sequence small RNAs, we employed ExtendAlign to pinpoint the identity percent of all sequences classified as bovine-specific with respect to human (Figure 6B). As expected, there is an ample percentage range at which the bovine-specific sequences redistribute, with the highest abundance peak at 75–77.5%; only a minor proportion was not aligned at all (Figure 6B, NA) —which would be expected only for those sequences accurately classified as bovine-specific. However, we find that ∼35% redistributed at 80–98% similarity range (Figure 6B, grey area). Hence, this suggests that a substantial amount of the bovine-specific sequences might have been the result of a high false-positive discovery rate. For example, a bovine a 22mer RNA sequence with an 80% identity to human —that is, four mismatches to its reference— is not necessarily unique to bovine.

**Figure 6.**
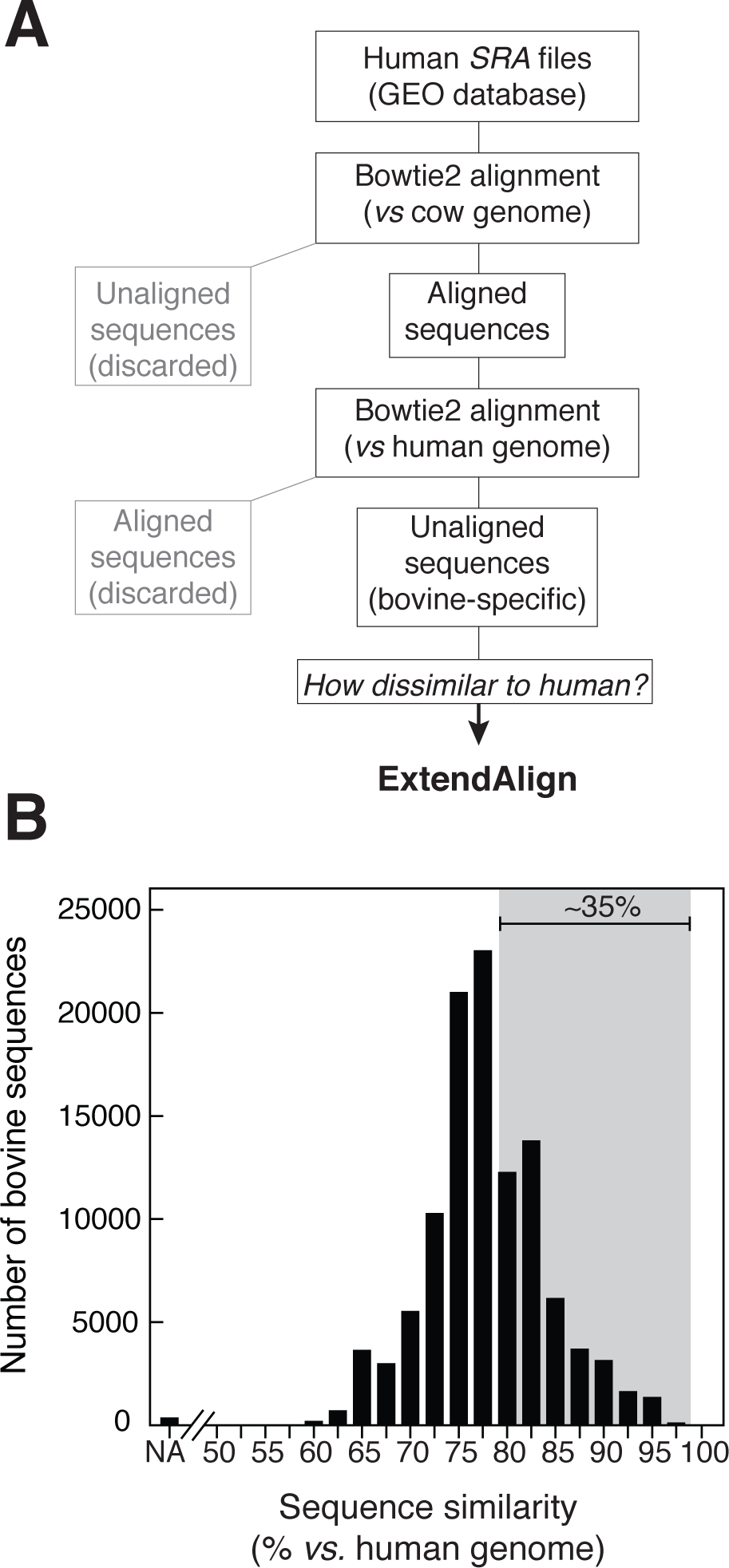
ExtendAlign pinpoints the similarity percentage of dissimilar short sequences. A, Diagram of pipeline followed to revisit the assignment of bovine-specific sequences from public databases with ExtendAlign. B, Histogram showing the abundance per sequence similarity as percentage of identity with the human genome. The grey area corresponds to sequences regarded as bovine-specific but have more than 75% similarity with the human genome. NA, not aligned.

## DISCUSSION

There has been a substantial effort by many research groups that focused on the development of aligners that reduce the processing time consumed for comparisons between sequences, which has yielded several reliable algorithms that perform robustly in the genomic era [47]. Most tools, however, have focused on improving the recognition of highly similar sequences to find alignments faster for homology and functional studies (27–29,40,45). In parallel to sequencing technologies that have increased the length of sequencing reads, most aligners also have adapted to handle longer sequences with the advantage of being also memory-efficient [4,30]. As a result, the alignments for dissimilar short sequences have been left somewhat unattended, and most algorithms have difficulties in providing accurate alignments for them. In this work, we show that most popular aligners handle perfect- or near-perfect match alignments well, which is not concerning when the interest is to align or map short sequences to a related reference, *e.g*., human small RNAs to the human genome [12]. However, this situation changes drastically when the similarity between sequences decreases. Although some aligners improved their results by modifying their parameters, others do not retrieve alignment results at all when presented with dissimilar short sequences. For those alignments that do take place, we found that aligners based on local algorithms have errors and tend to miss nucleotides at either 5’ or 3’ ends at regions that flank the seed substring. In contrast, global algorithms do not miss nucleotides but introduce gaps interspersed along the alignment to achieve end-to-end alignments —a useful feature to find *indels* [22,49]. Besides, global algorithms are impractical for finding imperfect alignments for sequences that differ in size, as they do not perform multiple hit alignments to a reference (Suppl. Figure S3A); this limits their utility to alignments with relationships already inferred. Thus, the problem of obtaining accurate reports for the alignment of short sequences has persisted over the years.

ExtendAlign is a post-analysis tool that identifies and corrects the errors originated by the alignment of dissimilar short sequences. It incorporates the tabular output of a local aligner and provides a list of all hits with their corrected m/mm for the total length of each query sequence. ExtendAlign improves the alignments by i) extending local alignments in an end-to-end manner; ii) including all nucleotides flanking the seed substring and therefore always fulfills 100% query coverage; iii) not increasing the total alignment size, nor introducing gaps artificially; and iv), increasing the sensitivity and specificity (Table 2). Thus, the results provided in the post-analysis lie at the intersection between those of local and global algorithms. Other algorithms combine local and global approaches (*i.e*., *Glocal*), but their recommended use and functionality is constrained to sequences longer than 100 nt (data not shown), eliminating its utility for the alignment of small RNAs [50]. In contrast, the recommended minimum size of the query of ExtendAlign is 8 nt, if primed with HSe-BLASTn (Suppl. Table S1).

We employed BLASTn as priming base because it provides the highest number of hits compared to the other aligners tested in this work, but we envision ExtendAlign could be adapted to be compatible with other local aligners too. Although ExtendAlign was initially developed to satisfy the need for multiple-hits pairwise alignments in an end-to-end manner of short sequences, it executes robustly with longer sequences, like precursor microRNAs, which are instruments frequently used for the discovery of novel microRNAs.

Since ExtendAlign is not an aligner itself, its recovery rate will match that of its priming base. For this reason, it will only correct alignments that took place by the aligner. Selecting HSe-BLASTn as priming base allowed to correct alignments resulting because of the high recovery rate, which also helped to pinpoint m/mm in alignments for short sequences with outstanding specificity and sensitivity that may otherwise go unnoticed by using other commonly employed aligners alone. Since full-length and perfect-match alignments are not prone to improvement, ExtendAlign is more robust in correcting the number of m/mm in more dissimilar alignments —a useful feature for alignments between evolutionarily distant species. Particularly, ExtendAlign refines the alignments of dissimilar short sequences situated within the twilight-zone, the identity percent barrier that impedes to estimate conservation reliably [25,26,46]. Since lncRNAs are subject to weak functional constraint and rapid turnover during evolution [51], their similarity percent spans across this range too [26], and therefore the use of ExtendAlign may aid in the study of their conservation and functionality if long sequences are aligned in shorter segments to a reference.

The intriguing finding that RNAs from bovine origin may transfer into cell cultures as a result of incubation with FBS has raised a controversy in the field of research of microRNAs [27,28]. Here we present evidence that at least some of the sequences considered bovine-specific by Wei and colleagues might be the result of a high false-positive discovery rate. Since those bovine-specific sequences were obtained with Bowtie2, the fact that upon a detailed analysis ∼35% of them showed high identity when aligned to human, validates the use of ExtendAlign to report the sequence identity for short sequences accurately. As no other tool combines the high accuracy and discovery rate of ExtendAlign in identifying m/mm for the alignment of short sequences, its use reduces the false discovery rate importantly. We anticipate that future studies will tackle with much better precision how this problem truly affects the entire literature in the microRNA field.

Since aligners are essential in computational biology, we anticipate that the utility of ExtendAlign can broaden up to other areas affected by the same type of bias in short sequence alignments, like the discovery of novel microRNAs [52,53], tRNA-derived fragments expression regulation [54], and other small RNAs from emerging model organisms [11]; to study homology of lncRNAs [55]; or to pinpoint similarity among pathogenic or complex samples (*e.g*., host-pathogen metagenomics) [56]; to name a few. The sequencing era has allowed us to analyze several species at the genomic level; the prediction of conserved and non-conserved microRNAs among species has become a field of great interest in the last decade. Thus, ExtendAlign can help in reducing false-positives commonly found in homology searches for small RNAs and also to increase the refinement of alignments between dissimilar sequences when high precision is needed.

## METHODS

### Datasets

Simulations were performed by building 8,500 random 22 nt long query sequences following a uniform probability distribution to model nucleotide content. The subjects were generated by a fixed seed of 7 nt for every sequence, with a start position chosen at random. One, or up to fifteen-nucleotide changes were introduced into the region corresponding to each query sequence. One insertion or deletion (indel) was introduced per every nine mismatches. The length of indels followed an exponential distribution with lambda = 2. Finally, randomly generated nucleotides were added to both ends of the previously mutated query sequences to yield a total length that ranged from 50 to 170 nt.

The datasets of mature and precursor microRNAs used as queries and subjects were downloaded from miRBase (release 22) [57,58]. The mouse (*Mus musculus*, mm10p6), armadillo (*Dasypus novemcinctus*, v.3.0), and platypus (*Ornithorhynchus anatinus*, v.5.0.1) genomes used as long-sequence subjects were downloaded from NCBI. The SRR515903 dataset for bovine-specific sequence analysis was downloaded from the Gene Expression Omnibus [27,59].

### HSe-BLASTn setup

BLASTn v2.8.1 [18,39] was used to prime and implement ExtendAlign seeds (Table 1). The parameters of the high sensitivity version of BLASTn, referred to as HSe-BLASTn in this work are as follows: word size: 7; reward: 1; penalty: -1; gap open: 2; gap extend: 2; e-value: 10; DUST: false; soft masking: false. The DUST filtering and soft masking options were disabled in order to keep all query sequences, even when scored as low complexity. HSe-BLASTn databases for subject sequences were built using the makeblastdb command line tool including the parse_seqids option (to keep the original sequence identifiers) and the dbtype nucl (specific for input nucleotide sequences) parameter.

### ExtendAlign Implementation

ExtendAlign was developed as a sub-modular project wrapped in a Nextflow environment [29]. At its core, multiple sub-modules combine in-house scripts (one for each pipeline stage), and implementations of BLAST+ v2.8.1 [18,39], bedtools v2.27.0 [60] and SeqKit v0.10.1 [61]. The sub-module scripting follows the mk syntax to establish input-output file dependency control [62]. ExtendAlign receives DNA/RNA sequences for query and subject in FASTA or multi-FASTA file format. The algorithm runs through the following general stages:

1. FASTA formatting. Sequence length is appended to FASTA headers.
2. Construction of BLAST database. Subject FASTA file is used to produce a BLAST database.
3. HSe-BLASTn alignment. Queries are aligned against subjects using the high sensitivity parameters of the HSe-BLASTn setup.
4. Hit filtering. HSe-BLASTn may have reported many alignment hits per query. At this stage, ExtendAlign provides the option to keep “all-hits,” or to find the “best-hit.” Best-hit was defined as the longest alignment with the least mismatches (including the query mismatches in the gaps).
5. Extension coordinates calculation. For each HSe-BLASTn hit, the length of the query and subject extensions is calculated as follows:

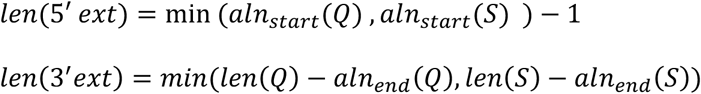

where *Q* and *S* refer to the query and subject sequences, respectively; *aln*_*start*_ and *aln*_*end*_ refer to HSe-BLASTn alignment start and end position, respectively; and *len* refers to the sequence length. The 5’ and 3’ extension regions are described by the vector defined by their start and end coordinates, determined by:

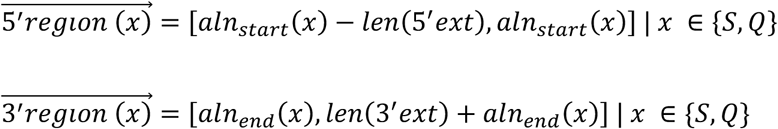 This pipeline stage yields a set of query and subject sequence coordinates from which nucleotides will be extracted based on each HSe-BLASTn hit. The outlined procedure self-adjusts when dealing with minus strand hits to enable ExtendAlign to work with any strandness configuration of a BLASTn run.
6. Extraction of extended nucleotides. Using the coordinates from the previous step and the original FASTA inputs for each HSe-BLASTn hit, the nucleotide sequences are appended at the query and subject 5’ and 3’ ends.
7. Percent identity recalculation. To allow RNA vs DNA comparison extended “U” nucleotides are transformed to “T”. Then, the corresponding (5’ *vs*. 5’ and 3’ *vs*. 3’) extended nucleotides are compared positionally to calculate extension mismatches; no gaps are allowed. The amount of the query covered by the HSe-BLASTn alignment and the extension phase (effective length = *eff*_*len*_) is calculated as follows:

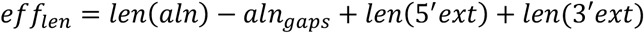

where *len(aln)* refers to the length of the alignment reported by HSe-BLASTn, and *aln*_*gaps*_ is the number of gaps introduced into the query sequence by HSe-BLASTn. The total number of mismatches found in the query is calculated as follows:

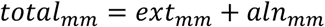

where *ext*_*mm*_ is the number of mismatches introduced during the extension phase and *aln*_*mm*_ is the number of mismatches reported by HSe-BLASTn. The identity percent calculation depends on the difference between the effective length and the query length:

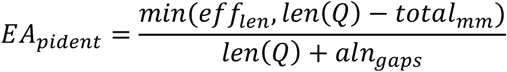
8. No-hit query appendage. Query names for which HSe-BLASTn did not find hits are appended to the correction table results as “NO_HIT” in the subject field.
9. Generation of alignment report. Results are gathered in a tab-separated file that reports all hits with a list of query and subject names, with percent identity before and after the ExtendAlign correction. NO_HIT queries are included in the report.

### Performance tests

BWA-MEM, BWA [32], BLASTn [18], BLASTn-short[33,39], Bowtie2 (end-to-end) [31,63] and ExtendAlign (all-hits mode) were used to align every simulated query sequence against the subject database. For Needle [22], we designed a computing cycle to align each query sequence against its corresponding subject sequence. Each algorithm was run twice, first with default parameters, then with permissive parameters. Permissive parameters were chosen to maximize the number of alignments to increase the chance of finding the true positive one. Alignments were considered true positive when queries aligned with their respective subjects and when the position of the fixed seed region (established during the generation of the data) matched the expected seed according to our simulated databases (see Datasets above). Recovery percentage was defined as the number of true positive alignments versus the total number of query sequences. The modified parameters for each algorithm are listed in the Suppl. Information. The information contains the corresponding command line used. Each nucleotide from the true positive alignment was assigned to one of four categories: true positive (TP), when the simulation and the aligner agreed in the existence of a change; true negative (TN), when the simulation and the aligner agreed in the absence of a change; false-positive (FP), when the aligner found a change that was not recorded in the simulation; and false-negative (FN), when the aligner did not find a change that was introduced by the simulation. Query nucleotides not included in the alignment span were considered as FN. Subject overhangs were considered as FP. Specificity was calculated as TN / (TN + FP), recall was calculated as TP / (TP + FN) and precision was calculated as TP / (TP + FP).

### Bovine-specific dissimilarity percentage calculation

The SRR515903 was converted into a FASTQ file with the SRA toolkit (v. 2.9.1). Bovine-specific sequences were extracted by aligning the library to cow (*Bos taurus*, bosTau8, UCSC Genome Browser) and human (*Homo sapiens*, Hg38p12, NCBI) as reference genomes with Bowtie2 [31,63], using the parameters defined in Wei *et al*. [27]: mode, local; seed length, 25; mismatches in seed, 0. Queries that aligned to the cow genome, and did not to the human genome were extracted into a new FASTA file for downstream analysis with ExtendAlign.

### Statistical Analysis

Data were plotted with no bins as cumulative frequency distributions. The significance between datasets was obtained by using the Kolmogorov-Smirnov test for non-parametric data with a significance value of *p* < 0.005. Non-linear fit curve by least squares is shown for every dataset with the coefficient of determination as a measure of goodness of fit. Box plots are presented as min to max values shown by quartiles; significance was obtained by using the Wilcoxon test for paired, non-parametric data with a significance value of *p* < 0.05. Simulated data was plotted with the R software. All other plots were done using GraphPad Prism software (v.7).

## DECLARATIONS

### ETHICS APPROVAL AND CONSENT TO PARTICIPATE

Not applicable

### CONSENT FOR PUBLICATION

Not applicable

### AVAILABILITY OF DATA AND MATERIAL

The datasets generated and/or analyzed during the current study are available in the Flores-JassoLab Github repository: https://github.com/Flores-JassoLab/ExtendAlign.

### AUTHORS’ CONTRIBUTIONS

MFT, JIHH, and IAO, constructed ExtendAlign and set GitHub repository; MFT, LGR, HT, SEAV and CFFJ, validated ExtendAlign; MFT, SEAV and CFFJ, envisioned the project and MFT, LGR, HT, SEAV and CFFJ wrote the manuscript. All authors read and approved the final manuscript.

## ACKNOWLEDGEMENTS

We thank all members of the Avendaño-Vázquez and Flores-Jasso laboratories for critical comments on the manuscript. We also thank Wayne Matten (NCBI-BLAST crew) for valuable help for implementing the command line HSe-BLASTn.

## TABLE LEGENDS

**Table 1.** Alignment performance of commonly used aligners for simulated short sequences.

**Table 2.** Alignment performance of short sequences with ExtendAlign.

