## Supplementary material for "ExtendAlign: the post-analysis tool to correct and improve the alignment of dissimilar short sequences": Suppl. Table and Figure legends

### ADDITIONAL FILE 1

### SUPPLEMENTARY TABLES

**Table S1.** Recommended use and features of aligners commonly employed for the alignment of short sequences.

**Table S2.** Number of total m/mm corrected by ExtendAlign.

### SUPPLEMENTARY FIGURE LEGENDS

**Figure S1.** Sequence dissimilarity impacts the alignment of short sequences. A, Two identical and two dissimilar short sequences were aligned with the aligners described. Red crosses indicate no alignment results. NA, not applicable. B, Aligners with hits under default parameters. C, Aligners with hits under permissive parameters. The report of each alignment is shown on the right. Bold red letters, nucleotides unreported; underlined letters, gaps reported.

**Figure S2.** Examples of biases generated by the alignment of dissimilar short sequences by global algorithms. A, The report by Needle considers the farthest 5′ and 3′ ends within the alignment, and overhanging positions are reported as gaps (underlined letters). B, Needle introduces gaps interspersed to achieve end-to-end alignments with a high recovery rate.

**Figure S3.** A, Local and global approaches retrieve different types of alignments for the same dissimilar sequences. HSe-BLASTn (grey) retrieves two different alignment hits to the same pair of sequences compared to Needle (cyan). B, Only the pairs of sequences with one-hit per alignment were presented to Needle from the total output of HSe-BLASTn.
