## Supplementary material for "ExtendAlign: the post-analysis tool to correct and improve the alignment of dissimilar short sequences": Suppl. Information

### SUPPLEMENTARY INFORMATION

### PARAMETERS FOR THE PERMISSIVE VERSIONS

Arguments modified for each tool used (according to each user manual).

**Bowtie (version 1.1.2)**

| | | |
|---|---|---|
| -v | 3 | Reports alignments with at most INT mismatches. Values allowed goes from 0 to 3. |
| -a | | Reports all valid alignments per read or pair (default: off). Validity of alignments is determined by the alignment policy (combined effects of -n, -v, -l, and -e). |
| --tryhard | | Finds valid alignments (trying as hard as possible) when they exist, including paired-end alignments. |

**Bowtie2 (version 2.2.6)**

| | | |
|---|---|---|
| --end-to-end | | Aligns the read from one end to the other, without any trimming (or "soft clipping") of characters from either end. |
| --score-min | L,-100,-1 | Sets the minimum alignment score needed for an alignment to be considered "valid". It is a function of read length. |
| -a | | Unlimits the number of alignments reported. |
| -L | 7 | Sets the length of the seed substrings to align during multiseed alignment. |
| -N | 0 | Sets the number of mismatches allowed in a seed alignment during multiseed alignment. |

**BWA-MEM (version 0.7.12-r1039)**

| | | |
|---|---|---|
| -L | 100 | (Clipping penalty). Keeps track of the best score reaching the end of query. Clipping will not be applied if it is larger than the best SW score minus the clipping penalty. |
| -k | 7 | Establishes the minimum seed length, matches shorter than INT will be missed. |
| -c | 500000 | Discards a MEM if it has more than INT occurrence in the genome. |
| -T | 0 | Discards alignments with score lower than INT. |
| -a | | Outputs all found alignments for single-end or unpaired paired-end reads. |

**BWA (version 0.7.12-r1039)**

| | | |
|---|---|---|
| -l | 7 | Takes the first INT subsequence as seed. |
| -n | 30 | Establishes the maximum edit distance, as the value INT. |
| -k | 0 | Establishes the maximum edit distance in the seed. |
| -M | 10000 | (Mismatch penalty). BWA will not search for hits with a score lower than (bestScore-misMsc). |
| -N | | (Disable iterative search). Finds all hits with no more than *maxDiff* differences. This mode is much slower than the default. |
| samse -n | 500000 | Establishes the maximum number of alignments to output. |

**BLASTn (version 2.2.31)**

| | | |
|---|---|---|
| -reward | 1 | Establishes the reward for a nucleotide match. |
| -penalty | -1 | Establishes the penalty for a nucleotide mismatch. |
| -strand | plus | Establishes the query strand(s) to search against database/subject. |
| -gapopen | 2 | Establishes the cost to open a gap. |
| -gapextend | 2 | Establishes the cost to extend a gap. |
| -word_size | 7 | Establishes the length of initial exact match. |
| -dust | none | Filters query sequence with “Dust”. |
| -soft_masking | false | Applies filtering locations as soft masks (i.e., only for finding initial matches). |
| -evalue | 5000 | Establishes the expected value (E) for saving hits. |
| -num_descriptions | 8500 | Shows one-line descriptions for this number of database sequences. |
| -num_alignments | 8500 | Shows alignments for this number of database sequences. |

**BLASTn-short (version 2.2.31)**

| | | |
|---|---|---|
| -reward | 1 | Establishes the reward for a nucleotide match. |
| -penalty | -1 | Establishes the penalty for a nucleotide mismatch. |
| -strand | plus | Establishes the query strand(s) to search against database/subject. |
| -gapopen | 2 | Establishes the cost to open a gap. |
| -gapextend | 2 | Establishes the cost to extend a gap. |
| -word_size | 7 | Establishes the length of initial exact match. |
| -dust | none | Filters query sequence with “Dust”. |
| -soft_masking | false | Applies filtering locations as soft masks (i.e., only for finding initial matches). |
| -evalue | 5000 | Establishes the expected value (E) for saving hits. |
| -num_descriptions | 8500 | Shows one-line descriptions for this number of database sequences. |
| -num_alignments | 8500 | Shows alignments for this number of database sequences. |

**ExtendAlign**

| | | |
|---|---|---|
| -strand | plus | Establishes the query strand(s) to search against database/subject. |
| -evalue | 5000 | Establishes the expected value (E) for saving hits. |
| -num_descriptions | 8500 | Shows one-line descriptions for this number of database sequences. |
| -num_alignments | 8500 | Shows alignments for this number of database sequences. |

### COMMAND LINE INSTRUCTIONS FOR DEFAULT VERSIONS

**Bowtie**

bowtie $INDEX_BOWTIE -f query.fa -S output.sam

**Bowtie2**

bowtie2 --end-to-end -x $INDEX_BOWTIE2 -f query.fa -S output.sam

**BWA**

bwa aln $INDEX_BWA query.fa >  output.sai
bwa samse $INDEX_BWA output.sai  query.fa >  output.sam

**BWA-MEM**

bwa mem $INDEX_BWA query.fa >  output.sam

**BLASTn**

```
blastn -query query.fa -db $DB_BLAST -task blastn -outfmt "6 qseqid sseqid pident length
mismatch gaps qstart qend sstart send evalue bitscore sstrand"
```

**BLASTn-short**

```
blastn -query query.fa -db $DB_BLAST -task blastn-short -outfmt "6 qseqid sseqid pident length
mismatch gaps qstart qend sstart send evalue bitscore sstrand"
```

### COMMAND LINE INSTRUCTIONS FOR PERMISSIVE VERSIONS

**Bowtie**

```
bowtie -v 3 -a --tryhard $INDEX_BOWTIE -f query.fa -S output.sam
```

**Bowtie2**

```
bowtie2 --end-to-end --score-min L,-100,-1 -a -L 7 -N 0 -x $INDEX_BOWTIE2 -f query.fa -S
output.sam
```

**BWA**

```
bwa aln -l 7 -n 30 -k 0 $INDEX_BWA query.fa >  output.sai
bwa samse -n 500000 $INDEX_BWA output.sai  query.fa >  output.sam
```

**BWA-MEM**

```
bwa mem -L 100 -k 7 -c 500000 -T 0 -a $INDEX_BWA query.fa >  output.sam
```

**BLASTn**

```
blastn -query query.fa -db $DB_BLAST -task blastn -reward 1 -penalty -1 -strand plus -gapopen 2 -
gapextend 2 -word_size 7 -dust no -soft_masking false -evalue 5000 -num_descriptions 11900 -
num_alignments 11900 -outfmt "6 qseqid sseqid pident length mismatch gaps qstart qend sstart
send evalue bitscore sstrand"
```

**BLASTn-short**

```
blastn -query query.fa -db $DB_BLAST -task blastn-short -reward 1 -penalty -1 -strand plus -
gapopen 2 -gapextend 2 -word_size 7 -dust no -soft_masking false -evalue 5000 -num_descriptions
11900 -num_alignments 11900 -outfmt "6 qseqid sseqid pident length mismatch gaps qstart qend
sstart send evalue bitscore sstrand"
```
